# How Stimulus Statistics Affect The Receptive Fields of V1 Cells

**DOI:** 10.1101/2021.03.08.434507

**Authors:** Ali Almasi, Shi Hai Sun, Molis Yunzab, Young Jun Jung, Hamish Meffin, Michael R. Ibbotson

## Abstract

We studied the changes that neuronal RF models undergo when the statistics of the stimulus are changed from those of white Gaussian noise (WGN) to those of natural scenes (NS). Fitting the model to data estimates both a cascade of linear filters on the stimulus, as wells as the static nonlinearities that map the output of the filters to the neuronal spike rates. We found that cells respond differently to these two classes of stimuli, with mostly higher spike rates and shorter response latencies to NS than to WGN. The most striking finding was that NS resulted in RFs that had additional uncovered filters than did WGN. This finding was not an artefact of the higher spike rates but rather related to a change in coding. Our results reveal a greater extent of nonlinear processing in V1 neurons when stimulated using NS compared to WGN. Our findings indicate the existence of nonlinear mechanisms that endow V1 neurons with context-dependent transmission of visual information.

## 1 Introduction

Our understanding of sensory coding in the visual system is largely based on the stimulus-response characterization of neurons. Traditionally, a basic set of stimuli (e.g., bars or gratings), were used to parameterize neuronal responses in terms of a restricted choice of stimulus parameters (e.g., orientation, spatial frequency etc.). This analysis allowed the measurement of tuning functions. However, such techniques provide only partial understanding of the neuronal response function and are particularly limited when the processing is nonlinear.

Later methodological improvements enabled the recorded responses of neurons to be characterized using statistically richer stimuli (Chen, Han, Poo, & Dan, 2007; Ringach, Hawken, & Shapley, 2002; Touryan, Felsen, & Dan, 2005). Recently, appropriate mathematical tools for a comprehensive neuronal characterization under arbitrary stimulus regimes have emerged (Fitzgerald, Sincich, & Sharpee, 2011; Liu et al., 2016; Rapela, Felsen, Touryan, Mendel, & Grzywacz, 2010; T. Sharpee, Rust, & Bialek, 2004). A prime example is the probabilistic framework wherein model estimation is performed by maximizing the likelihood of the model given the recorded responses and stimuli I. M. Park, Archer, Priebe, and Pillow (2013); M. Park and Pillow (2011). An example of this framework, which we have adapted, is the nonlinear input model (NIM) (Almasi et al., 2020; McFarland, Cui, & Butts, 2013). In this case, characterization is achieved by estimating the parameters of a receptive field model. Typically, the model first applies a cascade of linear filters on the stimulus. In the second stage, static nonlinearities map the output of the linear filters to neuronal spike rates (e.g., the generalized linear model).

How do these characterizations depend on the choice of the stimulus? To answer this question, first it is essential to control for artifactual fits, because the fitting method may extract statistical regularity inherent in the stimulus, rather than the stimulus-response relationship. To overcome this problem, maximum likelihood estimation can be used, as it provides an unbiased and consistent estimation method. Second, it is possible that the system may adapt to the stimulus statistics such that the stimulus-response relationship is altered. In the case of adaptation, it is the system that changes due to the relative presence of different types of features. Third, stimuli with different statistics may simply allow us to sample different operational regimes (Butts, 2019). For example, some regimes of operation are effectively unobservable or cannot be estimated because there are insufficient stimuli containing particular features to allow reliable estimation. In this study, our experiments aim to explore the last two points.

Here, we studied the changes that the NIM of cortical receptive fields (RF) undergoes when presented with different image statistics. We applied the NIM to the recordings of single cells in cat primary visual cortex (Fig. 1a). Cortical cells were stimulated with two types of stimuli which had distinct statistical properties: white Gaussian noise (WGN) and natural scenes (NS), with the same global Root-Mean-Square (RMS) contrast. WGN has a Gaussian distribution of contrasts, with a heavy over-emphasis on low contrasts. NS have more high contrasts and tend to be sparser. The NIM framework makes minimal assumptions about the kind of underlying neuronal processing and can fit the RF filters as well as a diverse range of nonlinearities that neurons employ to pool the output of their spatial filters (Almasi et al., 2020; McFarland et al., 2013). We estimated for each cell the spatial filters constituting the neuronal RF, and their corresponding pooling mechanism. The number of spatial filters for each cell is determined by a validation technique over a test dataset.

**Figure 1.**
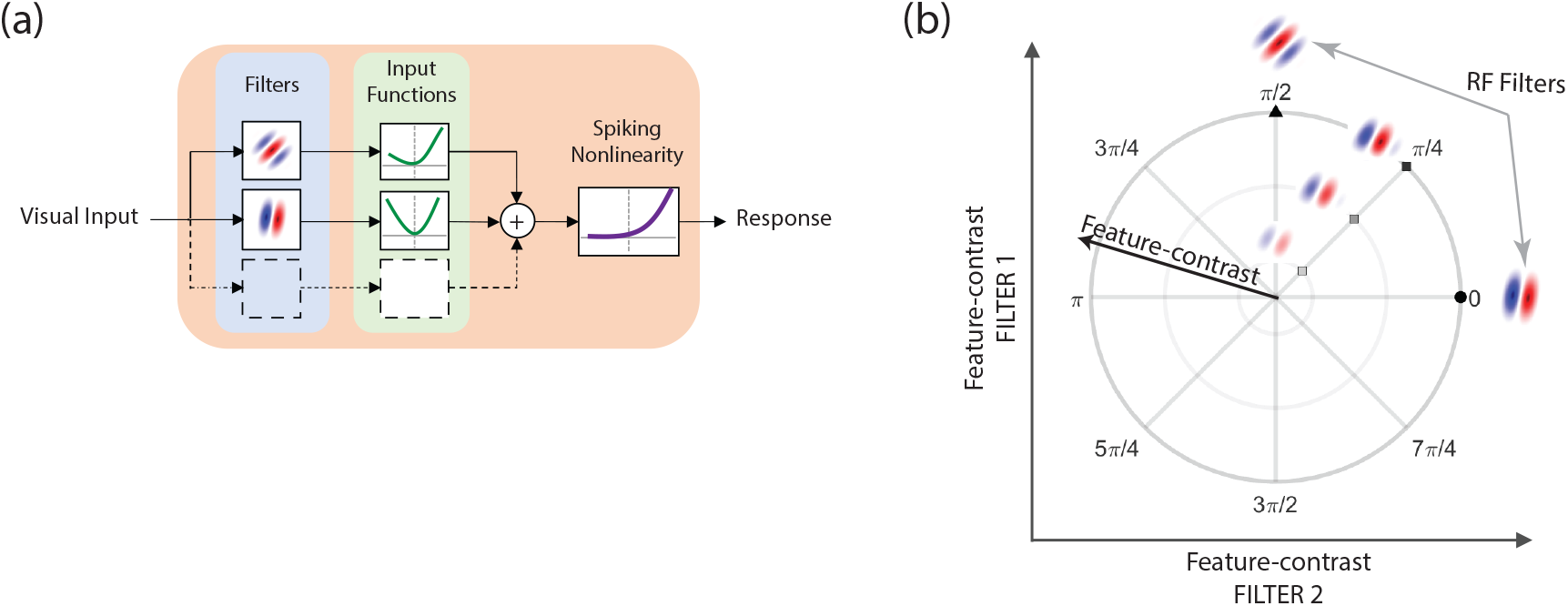
(**a**) Schematic diagram of the nonlinear input model (NIM) used in this study to characterize the spatial structure of V1 receptive fields. (**b**) Diagram depicts the concept of the feature space of a cell using its RF filters. The feature space contains any linear weighted sum of the RF filters.

We found that cells in primary visual cortex respond differently to the two stimulus types, with mostly higher spike rates and shorter response latencies to NS than to WGN. NS resulted in around twice as many uncovered RF filters compared to WGN. This difference was not related to the higher spike rates of cells to NS. Instead, we found that specific feature-contrasts attain much higher values in NS compared to WGN stimuli and are believed to be responsible for the differences in the number of RF filters. Our findings imply that cells in primary visual cortex adapt to the statistics of the visual stimulus to improve their coding efficiency, which enhances their capacity for information transmission.

## 2 Results

We studied the spatial structure of V1 receptive fields in anaesthetized cats identified using both White Gaussian Noise (WGN) and Natural Scenes (NS) as stimuli. The recorded neurons were visually stimulated with interleaved blocks of WGN and NS to increase the biological comparability of the recordings made with both stimuli (6 blocks of 12,000 stimuli per block for each stimulus type). The spatial RFs were uncovered using the NIM framework by maximizing the log-likelihood of the model given the pairs of stimuli and responses, which was done over the set of all model parameters simultaneously. The temporal RF of cells was left out in the modelling framework as (1) we wanted to investigate the changes that are brought about in the visual feature selectivity of cells, and (2) the used stimuli did not have enough temporal correlation to allow decent estimation of the temporal RF.

### 2.1 Differences between the V1 response profiles to WGN and NS

Although we did not study here the temporal aspects of the neuronal RFs, we observed major differences in the way that neurons responded to WGN and NS stimuli in terms of their response strength and latency (see Fig. 2a). Generally, V1 cells responded more strongly to NS stimuli (roughly 78% higher spike rates) than to WGN (Fig. 2b; mean ± standard deviation: 7.8±4.9 ips versus 4.4±3.3 ips). Furthermore, all cells showed longer response latencies to WGN than to NS stimuli, with an average difference of 11.4±6.2 ms (Fig. 2c; mean ± standard deviation: WGN = 29.0±6.3 ms versus NS = 17.6±4.4 ms). All analysis was performed on spikes occurring in the 33 ms window immediately after the latent period. While the latency was different for WGN and NS, the analysis window remained the same. Figure A.1 shows all responses from the units analyzed in this project, revealing that the peak responses occurred within the 33 ms window.

**Figure 2.**
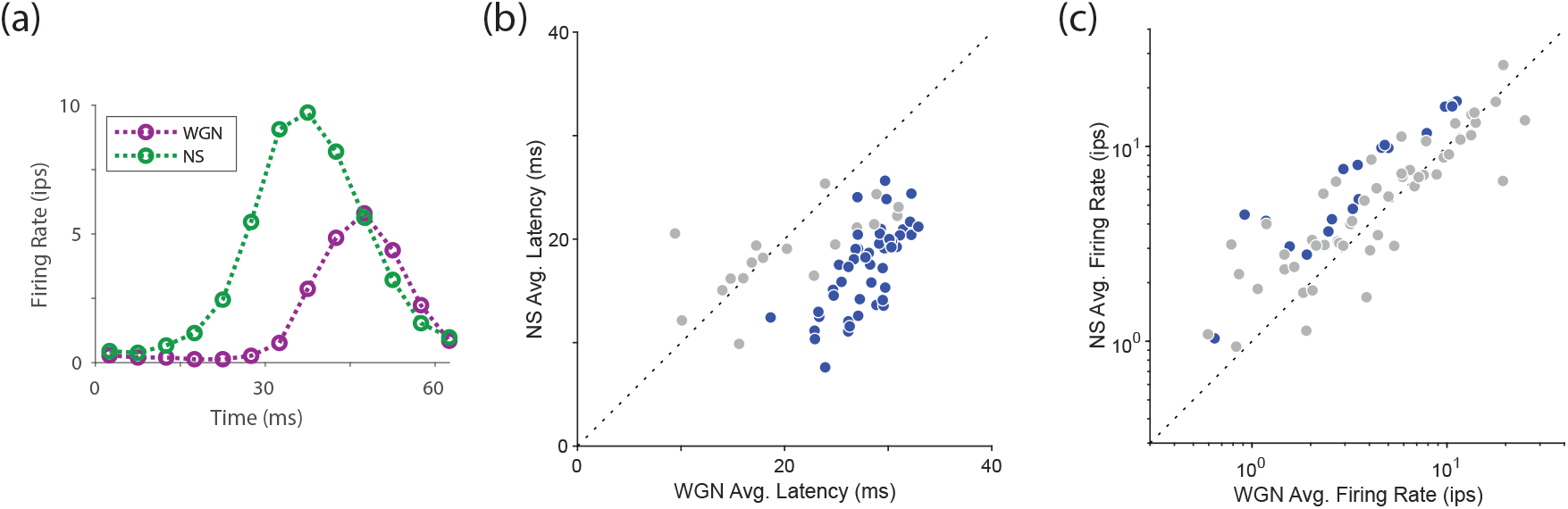
(**a**) Post-Stimulus-Time-Histogram (PSTH) of a V1 cell in response to WGN (magenta) and NS (green). There is a noticeable difference in the response delay between WGN and NS, with NS having faster rise time and therefore shorter delay. Scatter plots compare response profiles of V1 cells to WGN and NS stimuli in terms of (**b**) firing rate, and (**c**) response latency. Gray data points indicate cells for which the difference between WGN and NS was not statistically significant (unpaired t-test; *p >* 0.05). All cells that had significant difference between their WGN and NS response patterns showed higher firing rates and shorten response latency to NS than WGN. The plots show results of cells whose spatial RF structures were successfully characterized using both WGN and NS stimuli.

### 2.2 V1 RFs unveiled by NS are typically higher dimensional compared to those with WGN

We successfully uncovered spatial RFs for 92 orientation selective V1 cells using either NS or WGN (72 cells had RFs uncovered using both stimuli). Among these 92 cells, 87 cells had RFs uncovered using NS, 58 had RFs uncovered using WGN, and 53 cells had their RFs uncovered successfully using both stimulus types. In addition to the 92 cells with oriented RFs, we also uncovered RFs for 58 other cells whose RFs were non-oriented. According to a recent study (Sun et al., 2021), these cells might be of thalamic origin whose axons terminate in V1. Hence, since we were not certain about the cortical origin of these additional 58 cells with non-oriented RFs, we excluded them from our RF analysis.

Figure 3a presents the spatial RFs of an example V1 cell uncovered using the NIM framework under both NS and WGN. The most striking difference between these two model fits emerges in the number of spatial filters. Using the same number of stimulus images, the RF characterization using WGN and NS identified one versus three spatial filters, respectively. This was typical in our population of V1 cells, with a majority of cells (78%) having more spatial filters identified using NS than WGN (Fig. 3b-c). On the contrary, only a small fraction (8%) of cells had more uncovered spatial RF filters using WGN than NS. The distribution of the difference between the number of uncovered spatial filters using WGN and NS (#NS filters - #WGN filters) for the same cells varied from -1 to 4 (inset, Fig. 3c), but is asymmetric and heavily skewed to positive values. We investigate possible causes for having identified more spatial filters for RFs using NS than WGN in the following sections.

**Figure 3.**
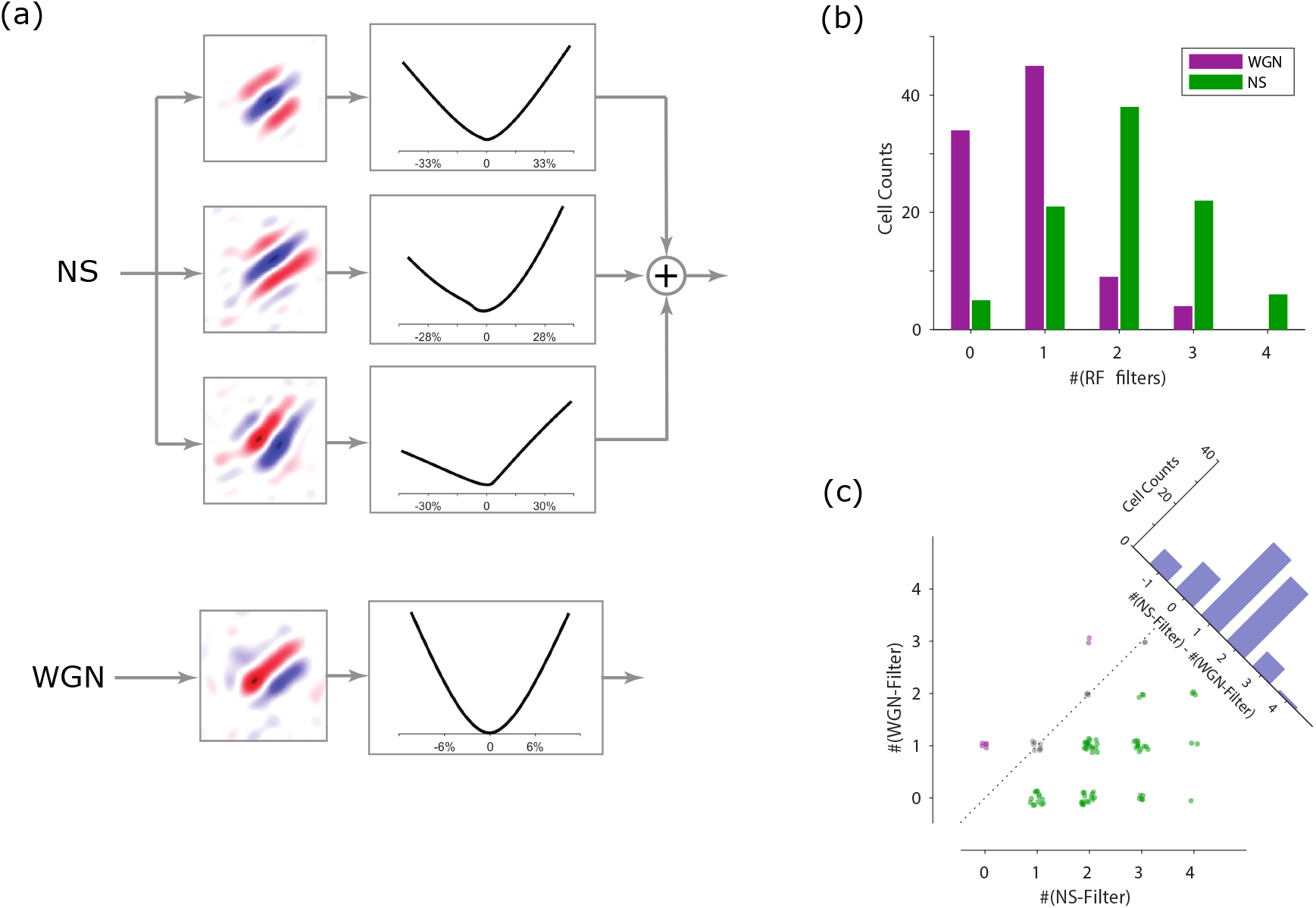
(**a**) Spatial RF structures of a V1 cell uncovered using the NIM and with NS (upper) and WGN (lower). For each model characterization only the spatial filters and input functions are shown as the spiking nonlinearity has a parametric log-exponential form. The abscissa in the Input Functions denotes the percentage of feature-contrast in terms of Michelson contrast. Red and blue in the filters indicate ON and OFF subregions of the filters, respectively. (**b**) Distribution of the number of spatial filters per each V1 cell, for the RFs uncovered using WGN and NS stimuli. (**c**) Scatter plot shows the number of RF filters uncovered using NS versus WGN stimuli, for each V1 cell. Data points are jittered as to provide a clearer view. The inset superimposed bar graph gives the distribution of the difference between the number of RF filters between NS and WGN.

### 2.3 Comparison of the feature spaces uncovered with WGN and NS

How does the identified spatial feature sensitivity of each cell depend on the type of stimulus (WGN or NS) used to estimate the model? For a given model, the set of spatial features to which the cell is sensitive is determined by its RF filters. The diagram in Figure 1a represents the RF model of a cell with multiple spatial filters. This cell is sensitive to any feature in an image corresponding to one of its RF filters, but also any feature that is a linear weighted sum of the RF filters. This is because such a summed feature would drive each RF filter and hence neural response. The set containing all possible linear weighted sums (i.e., linear combinations) of the RF filters is referred to as the cell’s feature space (see **Methods**). Figure 1b shows how we represent the feature space of the model cell. Different linear combinations of the features, corresponding to RF filters, are represented as distinct points in the feature space.

As most cells had at least as many RF filters uncovered using NS compared to WGN, we investigated whether the WGN feature space (H_*wgn*_) was a subspace of the NS feature space (H_*ns*_). In general, the WGN feature space can take any of the three illustrated forms in Figure 4a-c relative to the NS feature space. Namely, the WGN feature space (a) resides completely in the NS feature space, (b) is orthogonal to the NS feature space, (c) neither sits completely within nor is it orthogonal to the NS feature space. In the first case, the WGN feature space is a proper subspace of the NS feature space, while in the second case there is no overlap between WGN and NS feature spaces. In practice, even if the WGN feature space of a cell is essentially a subspace of its NS feature space, it is unlikely that the estimated feature spaces will precisely coincide due to noise in the model estimation process. In our data, initial estimates of feature spaces for all cells fell into the third category described above. However, it is necessary to determine whether the part of the WGN feature space that is not in the NS feature space is statistically significant. To do this, for each cell we first decomposed each WGN RF filter into two components. (i) A component that was the projection of the WGN RF filter onto the NS feature space, termed the projected component, which lies in the NS feature space ℍ_*ns*_ (black-colored vector; Fig. 4c). The space spanned by these projected components is termed the WGN-projected feature space denoted by 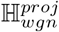. (ii) A component that is orthogonal to both the first component and ℍ_*ns*_, i.e., it sits outside the NS feature space ℍ_*ns*_, termed the residual component (gray-colored vector; Fig. 4c).

**Figure 4.**
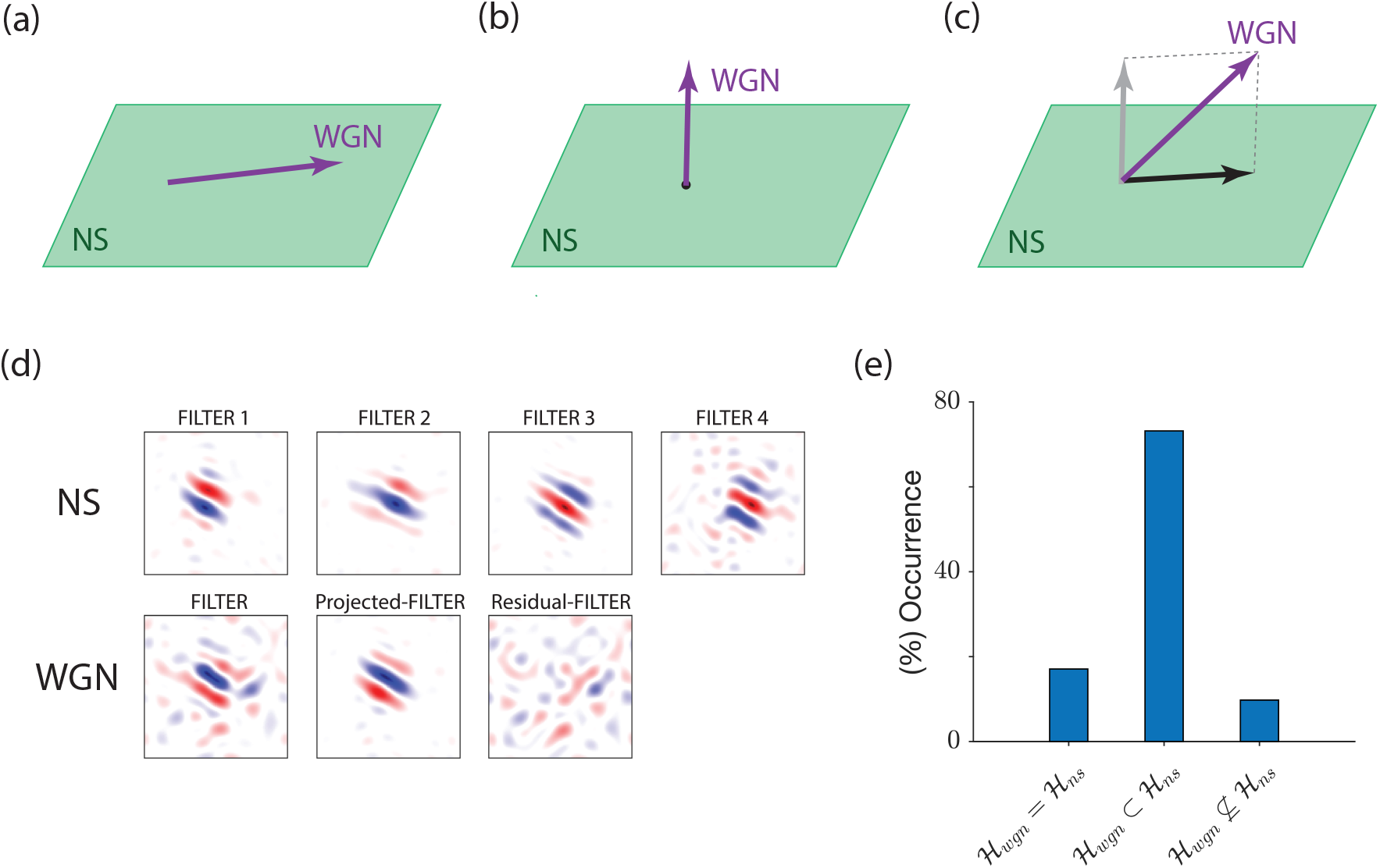
(**a**-**c**) The relationship between the uncovered WGN and NS feature spaces. (**d**) The first row shows the four RF filters of a V1 cell identified using NS. The second row from left to right shows the single RF filter of the same cell uncovered using WGN, its projection onto the NS feature space spanned by NS RF filters (shown in the first row), and its residual component that does not lie into the NS feature space. (**e**) Bar plot shows fractions of V1 cells for which WGN feature space is equal to, a proper subspace, or is not a subspace of the NS feature space.

Our visual inspection of the above components showed that in most cases the residual components of WGN RF filters, while non-zero, were noisy and had no meaningful structures (see

Fig. 4d). This suggests that they may not contribute significantly to model predictions. If this was so, then ℍ_*wgn*_ may be considered to be effectively a subspace of ℍ_*ns*_. To determine whether this was the case, we compared the predictive ability of the original WGN model with that of a model that was restricted to operate on only WGN-projected feature space, 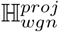; i.e., with the residual components removed (see **Methods**). After refitting the RF nonlinearities of this model on the WGN-projected feature space, we evaluated the performance of both models on a withheld WGN dataset. For a majority (90%) of cells in our V1 population the test showed an insignificant change in the model’s predictive ability, indicating that ℍ_*wgn*_ may be replaced by 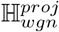, which is a subspace of ℍ_*ns*_ (Fig. 4e; unpaired t-test; *p >* 0.05). Hence, these cells had their ℍ_*wgn*_ equal to (17% of units), or a proper subspace (73% of units) of, ℍ_*ns*_. However, for the remaining minority (10%) of cells their response was predicted significantly less well by the new model using 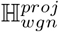, indicating that ℍ_*wgn*_ could not be considered to be a subspace of ℍ_*ns*_ in these cases.

### 2.4 Different dynamic range of V1 RF filters on NS and WGN

The spatial filter’s output quantifies the contrast level of the corresponding feature as it appears embedded in the WGN or NS stimuli, which is termed the feature-contrast (see **Methods**). It is a way of quantifying the contrast of those features within an image that drive a particular cell. While the overall RMS contrast of the WGN and NS was matched for our stimuli, the distribution of contrast of particular features could differ between WGN and NS due to their inherent statistical structure. For those features to which cells were sensitive, we often noticed considerable differences between the level of feature-contrast between WGN and NS. This is evident from Figure 5a, in which graphs show the distributions of feature-contrast of a typical V1 RF filter (inset) for WGN (magenta) and NS (green). The abscissa indicates the feature-contrast that is computed as the output of a normalized (unit norm) filter when applied on the stimuli. The presented distributions differ significantly in their spread. Defining a range of feature-contrast to span plus-and-minus three standard deviations (see **Methods**), the range of feature-contrast of the filter in Figure 5a on WGN and NS stimuli was measured as 1.7 and 5.9, respectively. It can be shown that these amounts correspond to 10% and 35% of the Michelson contrast of the feature.

**Figure 5.**
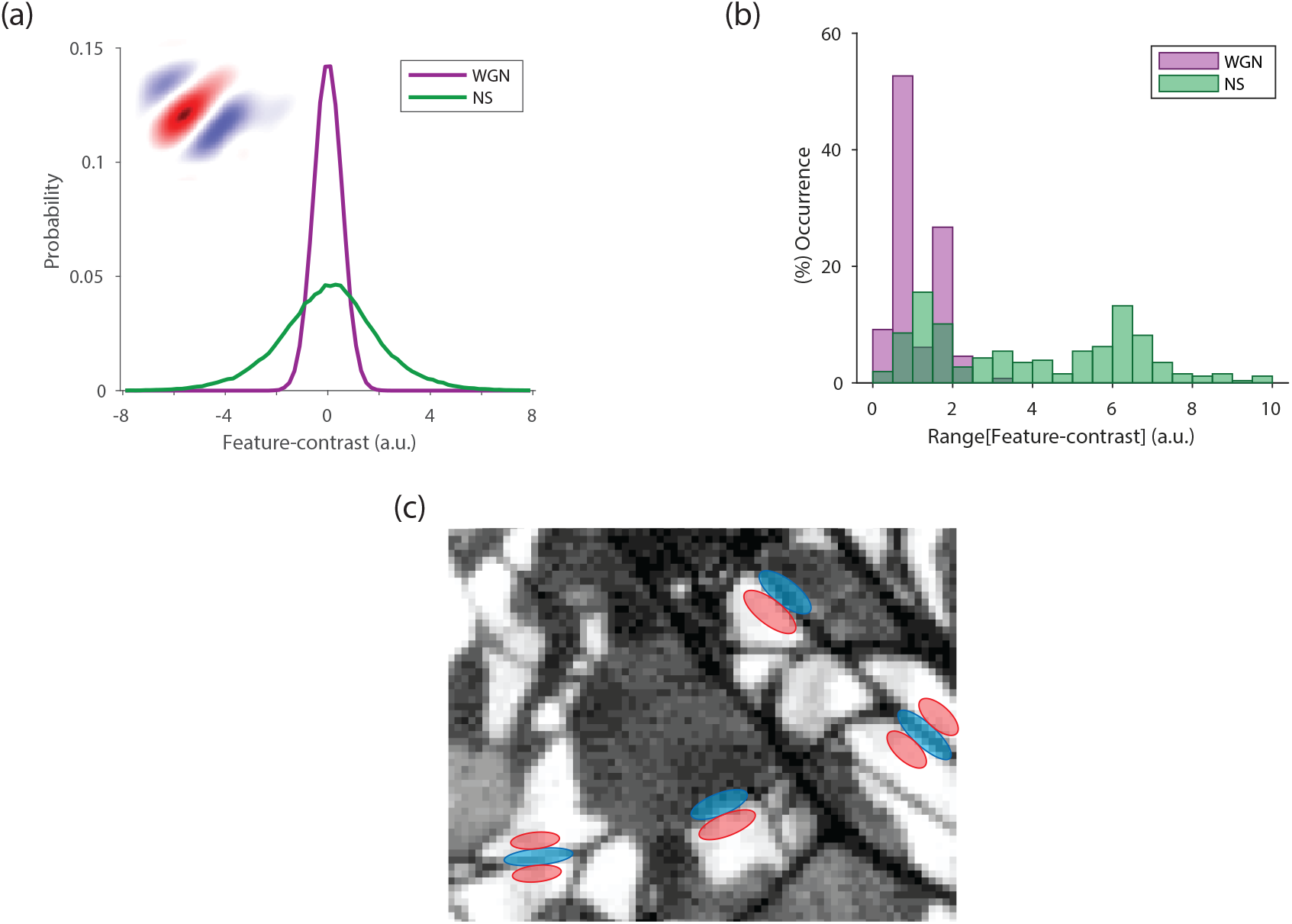
(**a**) The graphs show the empirical distribution of feature-contrasts of an example V1 RF filter (inset) obtained on WGN (magenta) and NS (green). (**b**) Bar graphs plot distributions of the range of the feature-contrast for the population of V1 RF filters on WGN (magenta) and NS (green). (**c**) Strong resemblance between the Gabor-like filters and the patterns that occur ubiquitously in NS indicate significantly larger values for the output of these filters (i.e., feature-contrast) on NS compared to WGN. Gabor filters are depicted as bands of red (ON) and blue (OFF) that are superimposed on parts of the scene that show a perfect match with their spatial structures.

The distribution of the range of feature-contrasts for the population of V1 RF filters uncovered using WGN and NS is given in Figure 5b, which demonstrates significant differences between these two stimulus types across the V1 population. Here, feature-contrast is computed as the output of normalized (unit norm) RF filters when applied on the stimuli. Such a pronounced difference arises due to the nature of the stimuli in relation to the feature sensitivity of V1 cortical neurons. As stated before, the differences between WGN and NS stimuli are in terms of second- and higher-order statistical dependencies between pixels, which are ubiquitous in NS but do not exist in WGN. The higher-order statistics of NS mainly account for structures like edges, curves and contours in these images. Of course, many V1 cells are highly responsive to these features because they have Gabor-like oriented RF filters. Accordingly, numerous occasions can occur in natural scenes in which a Gabor-like RF filter of a V1 cell is partially or highly matched with a feature in the scene (like the ones illustrated in Fig. 5c). This results in large values at the output stage of V1 RF filters, namely the filter’s feature-contrast. This provides an intuitive explanation for the higher range of feature-contrasts of V1 RF filters on NS than on WGN (Fig. 5b). Further, this considerable difference between the distributions of feature-contrast for V1 RF filters on WGN versus NS implies that the dynamic range of RF filter outputs that drive V1 cells is significantly larger when operating on NS than on WGN.

### 2.5 Difference in the feature-contrast response functions

We examined how each cell’s nonlinear response to feature-contrast depended on stimulus type (i.e., WGN vs NS). In the case of cells with multiple spatial filters, this feature-contrast response function determines how the cell pools those corresponding features in the visual input to give a response. For cells with equal numbers of RF filters between WGN and NS, we use the feature directions that are shared between the uncovered WGN and NS feature spaces to compare the response functions between the two stimulus regimes. The shared dimensions between the two feature spaces correspond to the WGN RF filters projected onto the NS feature subspace (i.e., 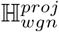). As we showed earlier, for most V1 cells (90%) the uncovered WGN feature space resides within the NS feature space. For cells that had more NS feature dimensions than WGN, we compare response functions on the feature directions that were shared between the two feature spaces, and also on NS feature directions that were orthogonal to the shared dimensions.

For simplicity, we begin by presenting the comparison of the response functions of cells that have a single RF filter using both WGN and NS. For these cells, 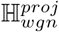 is the same as ℍ_*ns*_ (since there is only one filter dimension). For most cells, such as that illustrated in Figure 6, the WGN and NS feature-contrast response functions did not coincide and were significantly different (bottom Fig. 6b; filled symbols) (unpaired t-test; *p <* 0.001). Note that the comparison of feature-contrast response functions can only be made on the smaller shared domain of these functions, which corresponds to the range of feature-contrast for WGN (Fig. 6b). Cells often exhibited a change in their response functions between WGN and NS that were akin to the pattern shown in Figure 6b. Qualitatively, the changes were often characterized by an increase in spike rate in response to WGN over NS at the same low level of shared feature-contrast (bottom Fig. 6b). Note that despite this, the overall mean spike rate to NS was often higher than to WGN, presumably because the level of feature-contrast was higher for NS.

**Figure 6.**
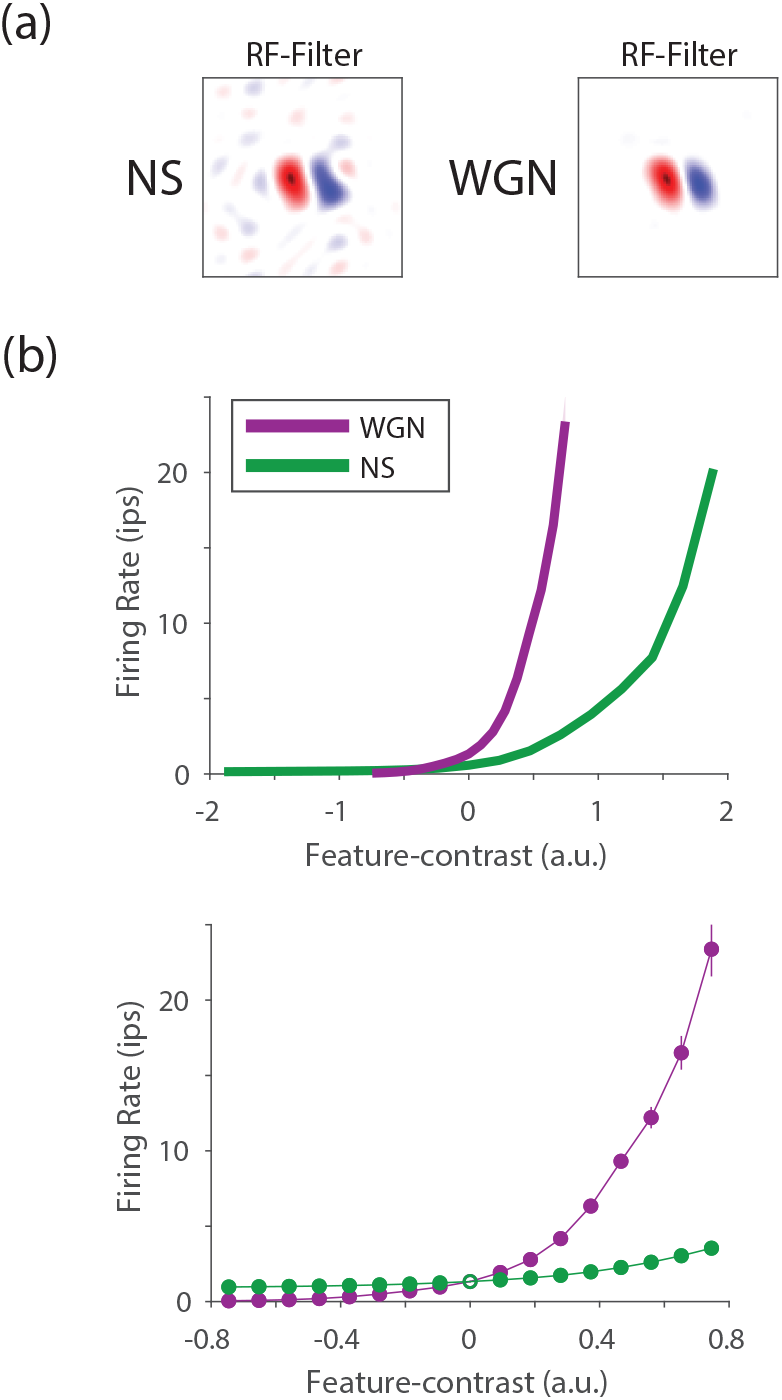
The changes observed in the feature-contrast response functions of V1 cells due to the change in the stimulus statistics from WGN and NS. (**a**) RFs filters estimated using the NIM and with NS (left) and WGN (right) stimulation for an example V1 cell. (**b**) Top panel compares Feature-contrast response functions identified using WGN (magenta) and NS (green) for the same V1 cell. Bottom panel compares the same functions but zooms into the range of WGN feature-contrast.

Figure 7a-c presents the response functions for a cell that had multiple but equal feature dimensions on both WGN and NS. Here, the comparison is similarly performed on the WGN-projected filter dimensions. Figure 7d-f shows the response functions for a cell that had more NS feature dimensions than WGN. In this case, the comparison is done on the feature dimensions that were shared between the two feature spaces (WGN-projected filter dimensions; Fig. 7e), and the NS feature dimensions that were orthogonal to the shared dimensions (NS-orthogonal dimensions; Fig. 7f).

**Figure 7.**
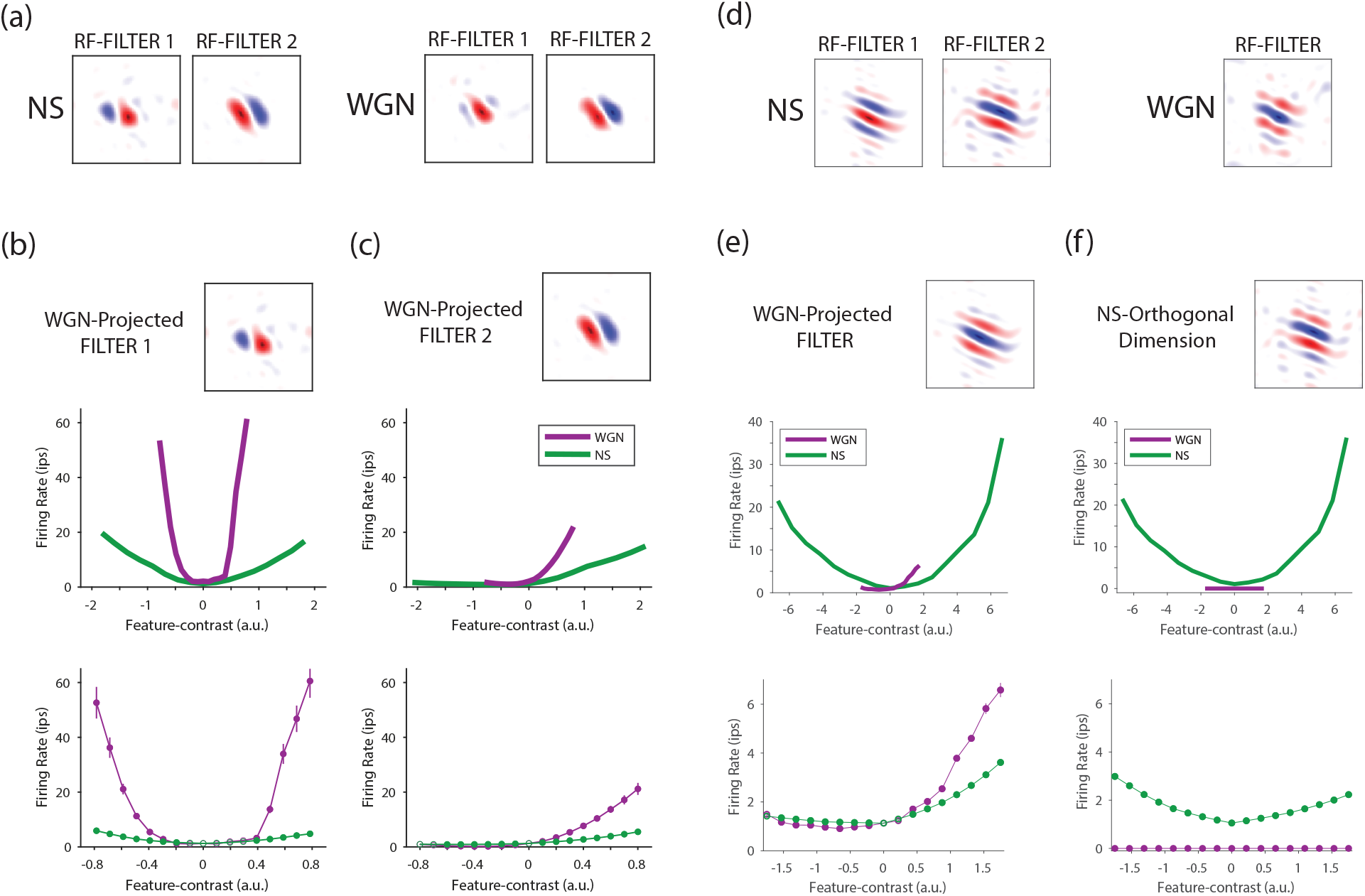
(**a**) RF of a V1 cell that resulted equal number of filters identified using the NIM and with NS (left) and WGN (right). Top panels in (**b**-**c**) compare the feature-contrast response functions of the same cell estimated using WGN and NS, upon the presented WGN-projected filters. Bottom panels compare the same functions but zoom into the range of WGN feature-contrast. (**d**-**e**) RF of a V1 cell that had two filters identified using NS but had one filter using WGN. Comparisons of the feature-contrast response functions are given in (**e**-**f**) with the same convention as in (**b**-**c**).

We quantified these changes using an index of normalized Difference between Area Under the Curve (nD-AUC) of the WGN and NS feature-contrast response functions (Fig. 8a). This index was calculated for the polarity of feature-contrast to which the cell showed the strongest response dependency. The difference between the area under the curve of WGN and NS response functions is depicted using the yellow highlighted area in Figure 8a, which indicates how different the WGN response function is from the NS response function within the WGN feature-contrast range (Σ_*wgn*_). To obtain a normalized index (nD-AUC) that varies between -1 and 1, we divided the yellow highlighted area by the grey-shaded area, which is determined by the WGN feature-contrast range (Σ_*wgn*_), the maximum response of the cell within this range to either WGN or NS (*R*_*max*_), and the cell’s response at zero feature-contrast (*R*_0_). We computed this normalized index for the cells whose feature-contrast response functions featured a significant difference between WGN and NS (unpaired t-test; *p <* 0.001). Positive and negative values of nD-AUC indicate that the WGN response function sits either significantly above or below the NS response functions, respectively, within the WGN feature-contrast range. Across our population of V1 cells, 52/53 cells (98%) showed a positive value of nD-AUC, indicating that for them the NS feature-contrast response functions sit below the WGN response functions across the shared (projected) feature directions between NS and WGN (Fig. 8b; blue bars). For only one cell, the change between the NS and WGN response functions was statistically indistinguishable (at *p* = 0.001 confidence level). For cells whose dimensionality of WGN feature space was less than that of NS feature space, we considered and compared the response nonlinearities on the shared dimensions as well as the NS feature dimensions that were orthogonal to the shared dimensions. In contrast to the shared feature dimension, for orthogonal feature dimensions the nD-AUC indices were negative (Fig. 8b; gray bar), which is related to the trivial (i.e., constant) dependency of WGN response functions. The distribution of the nD-AUC values of all feature dimensions for cells in our V1 population is given in Figure 8c. Most positive nD-AUC indices vary between 0.1 and 0.4, with a peak at 0.3. Most negative nD-AUC indices vary between -0.2 and -0.4.

**Figure 8.**
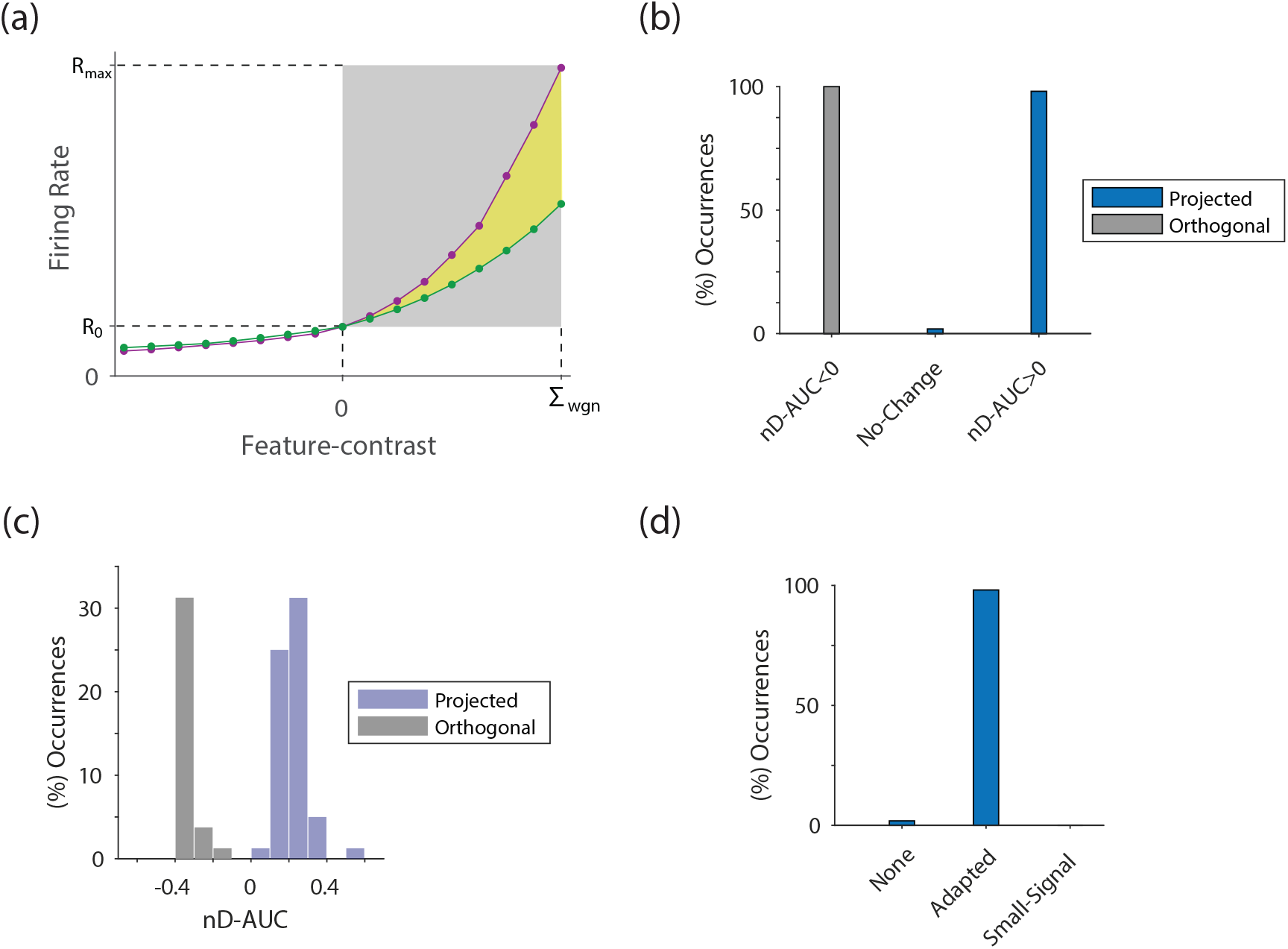
(**a**) Quantitative characterization of the difference between the feature-contrast response functions of an example cell uncovered with WGN (magenta) and NS (green) using the introduced index of normalized difference between the area under the two curves (yellow colored area). (**b**) Bar graphs present fraction of V1 cells whose feature-contrast response functions during WGN stimulation was significantly above (nD-AUC*>*0) or below (nD-AUC*<*0) their response functions during NS stimulation, upon feature dimensions that were shared (projected, blue bars) between the two feature spaces, or were orthogonal to the shared feature space (orthogonal, gray bars). No-change corresponds to those that did not show any significant changes in their feature-contrast response functions between WGN and NS. (**c**) Distribution of nD-AUC values of all feature dimensions. (**d**) The bar graph gives proportions of cells that showed no change, adaptation, or small signal effects.

### 2.6 Explaining the observed changes between the two stimulus regimes

For most V1 cells, the uncovered RF using NS stimuli reveals a larger repertoire of feature sensitivity compared to RF structure revealed with WGN (see Fig. 3a and Fig. 4d). The larger repertoire can be an indication of more complex feature selectivity and also the capacity for invariance (see Almasi et al. (2020)).

Here, we postulate different hypotheses that might explain the observed differences between the identified RFs of V1 cells using NS and WGN stimulus regimes.

#### 2.6.1 The observed changes are not artifactual nor methodological

One may suspect that the observed changes are methodologically related to our RF identification technique, because the fitting procedure may extract statistical regularities inherent in the stimulus, rather than the stimulus-response relationship. This has been long considered as a strong possibility for some RF identification techniques such as spike-triggered average and covariance when used in conjunction with statistically rich stimuli such as NS (Paninski, 2003; Schwartz, Pillow, Rust, & Simoncelli, 2006; T. Sharpee et al., 2004). However, this is not an issue here since we employed maximum likelihood estimation that is an unbiased and consistent estimation method, thereby, minimizing the possibility of artifactual RF filters obtained using NS (Paninski, 2004).

### 2.6.2 Greater number of RF filters using NS are not due to higher spike rates

We found that neurons in V1 were usually more responsive to NS than to WGN. One theory is that the larger number of uncovered RF filters with NS is related to the higher spike rates, because more spike data is available for the NIM fitting. The logic behind this suggestion is that having more spikes for fitting might lead to better fits, which might identify more significant filters. We assessed this hypothesis by performing a control analysis to understand the effect that spike count (number of spikes per stimulus image) might have on the number of uncovered spatial filters. In this analysis, for each cell we matched the cell’s spike count on NS and WGN (see **Methods**). It should be noted that this analysis was done for cells whose RFs were uncovered using both WGN and NS, but had more filters using NS (n = 32). After matching the cells’ spike counts between WGN and NS, we performed our statistical significance test (Z-Score *>* 2; see **Methods**) to determine the number of spatial filters within the RF with the matched (controlled) spike-count for NS. Most cells (83%) still had more filters within the RF identified using matched-spike-count-NS than using WGN (Fig. A.2), indicating that for most cells identifying a larger numbers of RF filters using NS is not attributed to there being more spikes for these stimuli compared to WGN.

As an additional check that our model fitting procedure correctly estimated the number of filters for WGN compared to NS, we refitted the model nonlinear functions to WGN, but provided the larger set of RF filters estimated for NS as fixed parameters in the model fitting. This test considered the possibility that our model fitting procedure might fail to estimate the full set of filters obtained for NS due to some flaws. If this was true, the refitted model with all the NS RF filters should have better predictive ability on a withheld WGN dataset, compared with the original WGN model with fewer RF filters. Nonetheless, our analysis proved that the models with the full set of NS RF filters never provide a better predictive ability than the simpler WGN models when tested on a withheld WGN test set (see **Supplemental Information** A). This supports the conclusion that a model fitted to NS data typically had a higher dimensional feature space compared to those fitted to WGN.

#### 2.6.3 The observed changes are not due to a small signal effect

Another possible explanation for the smaller number of RF filters typically found for WGN compared to NS could be a small signal effect. The argument in support of this effect is that stimuli with different statistics allow us to sample different operational regimes of the visual system, which might be effectively unobservable or cannot be estimated using some other stimulus types because there are insufficient stimuli containing particular features to allow reliable estimation. Recall from **Section** 2.4 that the range of feature-contrast with WGN was significantly smaller than that for NS. It could be that within the more restricted range of feature-contrast present in WGN that two models are not significantly different from each other; in this case for the extra dimensions of a model fitted to NS, the feature-contrast response functions should be statistically indistinguishable from the feature-contrast response functions of the model fitted to WGN, which is equal to zero in these dimensions. In this hypothesis, the additional filter dimensions present in the model fitted to NS are only able to be identified because the feature-contrast in NS becomes sufficiently large to identify a response that departs significantly from zero.

In our population of V1 data, we found no cell that showed any such small signal effect based on the above definition. For every cell where we found a reduction in the number of RF filters when the stimulus was changed from NS to WGN, we also found a significant change in their feature-contrast response functions within the restricted range of feature-contrast of WGN (Fig. 8d).

#### 2.6.4 Adaptation amongst other nonlinear phenomena to explain the changes

Based on our analyses, it appears that the changes in the RFs of V1 cells as a result of changes in the stimulus from NS to WGN are brought about using nonlinear phenomena. A possible explanation for these results may be the adaptation of cells to stimulus statistics. This is a potential explanation for the changes that we have observed in the RFs of V1 cells under WGN and NS stimulation. That said, our analyses cannot entirely rule out the possibility of other effects influencing the RFs, such as surround suppression, or end-stopping, which have been found for V1 cells when probing with basic visual stimuli such as bars and gratings.

In our V1 data, all cells underwent nonlinear effects in their response functions manifested as a significant amplification in their feature-contrast response functions across the restricted, shared feature dimensions when the stimulus changed from NS to WGN. However, across the feature dimensions that were orthogonal to the shared space (i.e., NS-orthogonal), WGN feature-contrast response functions exhibited trivial dependencies, resulting in a significant reduction in the response function when the stimulus was switched from NS to WGN (see Fig. 8c).

#### 2.6.5 Changes in the characteristics of RF filters

A thorough comparison between the WGN and NS feature spaces can be achieved by sampling features from these spaces and calculating the range of characteristics (e.g., orientation, spatial frequency, and spatial phase) of those sampled features (see Almasi et al. (2020)). However, this is a very significant project and is beyond the scope of the present paper. One issue that hinders direct comparison of the characteristics of RF filters between WGN and NS is the change in the number of RF filters. This can be particularly problematic when comparing the orientation characteristics between filters as we occasionally observed cells whose extra NS RF filters showed misalignment in their orientation preferences. To allow comparison with previous work (T. O. Sharpee et al., 2006), here, we will consider spatial frequency. We found that changes in spatial frequency were less problematic. We computed and compared the peak spatial frequency and spatial frequency bandwidth of filters within the same cell between the WGN and NS RFs. Preferred spatial frequency was preserved (Fig. 9a; r=0.58). The spatial frequency bandwidth showed a lesser degree of correlation (Fig. 9b; r=0.24). While the overwhelming finding for most cells was a similarity in spatial frequency tuning, at the population level there was a statistically significant bias towards filters with higher spatial frequency and narrower spatial frequency bandwidth during NS stimulation (unpaired t-test; *p <* 0.01).

**Figure 9.**
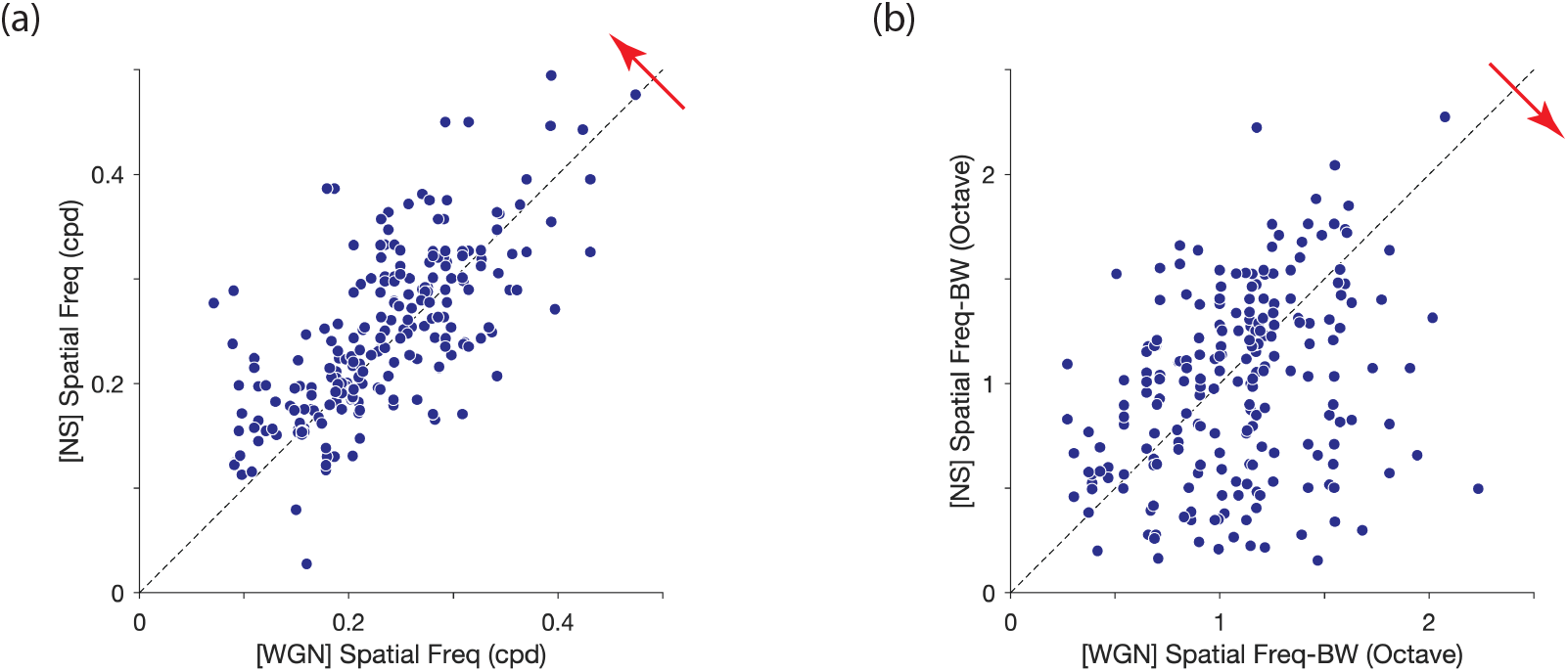
Scatter plots compare (**a**) spatial frequency preferences, (**b**) spatial frequency bandwidths of the RF filters uncovered using WGN and NS for each cell in our population of V1 cells. The mean of the distributions was found to be significantly biased towards the directions indicated by the red arrows.

## 3 Discussion

We studied the differences in the RF characteristics of cells in cat V1 when stimulated with white Gaussian noise (WGN) and natural scenes (NS). The RFs were estimated using a general class of neural model, the nonlinear Input Model (NIM), which allows characterization of RFs in terms of a cascaded linear-nonlinear architecture, with filters and nonlinear functions applied on the filters’ outputs. These outputs are referred to as feature-contrasts. For a majority of V1 cells, we found that the RF structure uncovered using NS resulted in more filters than did WGN. We also noted that the use of NS led to an increase in spike rate but the increase in filter numbers was not linked to the increase in spike rate. Rather, we believe it is attributed to the wider range of feature-contrasts generated by RF filters when using NS stimuli.

### 3.1 Contrast

The WGN and NS stimuli used in our experiments, though matched in global RMS contrast, are different in their local contrast. Small image patches (size 5*°*-7*°*) randomly sampled from NS show substantial variations in their RMS contrast, whereas the RMS contrast of patches sampled randomly from WGN are concentrated around the global contrast. The significant differences in the local contrasts of WGN and NS is related to different statistical regularities within these images (see Frazor and Geisler (2006)). WGN images lack any second- and higher-order statistics and are spatially stationary. That is, local statistics of random patches are independent of where in the scene the patches are sampled. Unlike WGN, NS have strong second- and higher-order statistics and are highly spatially non-stationary.

Our findings indicate that the larger range of feature-contrasts of V1 RF filters using NS, when compared to WGN, is due to the differences between the statistics of these stimuli, which is consequently related to the differences in their local contrasts. In this context, feature-contrast can serve as an indication of local contrasts in a scene.

### 3.2 Adaptation

We found significant changes between the way that V1 cells operate during WGN and NS stimulation. The differences in RF structures suggest the existence of nonlinear phenomena affecting V1 RFs, e.g., adaptation in the visual system, which may occur due to the changes in the statistics of the stimulus. Additional nonlinear phenomena such as surround suppression (Jones, Grieve, Wang, & Sillito, 2001; Webb, Dhruv, Solomon, Tailby, & Lennie, 2005; Wissig & Kohn, 2012) or end-stopping (Bolz & Gilbert, 1986; DeAngelis, Freeman, & Ohzawa, 1994) outside the classical RF may also contribute to the observed changes in our V1 data. Our RF characterization technique is capable of revealing such effects, given that the nature of the suppressive contribution is additive. However, it is possible that these effects are so subtle, or not localized enough to be identified using our RF characterization during NS and WGN stimulation.

Adaptation is often defined and characterized as a model-dependent phenomenon (Baccus & Meister, 2002). In general, any change in the parameters of the model describing the neuronal RFs, or any change in the description of the model is considered as an adaptation effect. In the latter case, the model is no longer capable of describing the neuronal responses to the original scene after adaptation to a new scene. As merely adjusting the parameters of our model when transitioning from WGN to NS, or vice versa, is inadequate for describing the neural data, our results reveal that a change in stimulus necessitates employing a different model.

A number of studies have investigated the adaptation caused by image statistics in the visual system (Felsen, Touryan, Han, & Dan, 2005; Lesica et al., 2007; Tkačik, Ghosh, Schneidman, & Segev, 2014). R. Shapley (1997) argued that contrast and scene statistics for a given spatiotemporal pattern are tightly related, suggesting that adaptation to scene statistics is equivalent to contrast adaptation. This is pertinent to the present study in which differences between the statistics of WGN and NS led to differences in the contrasts of the corresponding features in each cell’s RF filters, which accordingly altered the cell’s feature-contrast response function (see also Lesica et al. (2007)).

The effect of contrast adaptation has been widely studied at different stages in the visual system, from retina to primary visual cortex and higher cortical areas (Albrecht, Farrar, & Hamilton, 1984; Baccus & Meister, 2002; Bonin, Mante, & Carandini, 2006; Felsen et al., 2005; Kastner & Baccus, 2011; Lesica et al., 2007; Ohzawa, Sclar, & Freeman, 1985; Sceniak, Ringach, Hawken, & Shapley, 1999; R. Shapley, 1997; R. M. Shapley & Victor, 1978; Smirnakis, Berry, Warland, Bialek, & Meister, 1997; Solomon, Peirce, Dhruv, & Lennie, 2004; Zaghloul, Boahen, & Demb, 2005). The findings suggest intrinsic contrast adaptation effects at each stage of visual processing, which are inherited by later stages. Here, we will discuss possible mechanisms that can lead to our observed findings based on contrast adaptation effects in the visual system.

Numerous retinal studies in various species found that a change in contrast brings about a significant change in the temporal properties of retinal ganglion cell (RGC) responses (Sceniak et al., 1999; R. Shapley, 1997; R. M. Shapley & Victor, 1978; Smirnakis et al., 1997; Zaghloul et al., 2005). For a given image, the retina is forced to compromise between sensitivity and localization of light stimuli. Sensitivity is achieved using longer integration times for low contrast stimuli (R. Shapley, 1997), whereas localization is achieved using less surround integration for high contrast stimuli (Sceniak et al., 1999). At low contrasts, RGCs increase their synaptic integration time to improve sensitivity to the stimulus feature, leading to increases in response latency. This increased response delay then propagates throughout the visual hierarchy, thereby causing longer latencies for visual cortical cells, as noted in our data for WGN.

#### 3.2.1 Explaining changes in filter numbers

The dimensionality of the feature space of a cell directly relates to the complexity of its RF. RFs of simple cells often comprise a single filter (Almasi et al., 2020). A shift from multiple to single filters has been inferred from experiments with grating stimuli (Cloherty & Ibbotson, 2015; Crowder, Van Kleef, Dreher, & Ibbotson, 2007; Meffin, Hietanen, Cloherty, & Ibbotson, 2015). In those experiments, when contrast was reduced, cell responses shifted from those expected from RFs with multiple filters (complex-like) to those expected from a single filter (simple-like). A possible explanation for this switch might be adaptation to image contrast (or scene statistics). When presented with WGN, RGCs adapt to the low feature-contrasts of their RF filters. This adaptation introduces a change in the feature-contrast response functions of RGCs, known as contrast gain control (Smirnakis et al., 1997). This leads to a reduction in the contrast gain and consequently the threshold of the feature-contrast response function (Fig. 10). The effect is to increase the cells’ sensitivity to the low feature-contrast of the stimulus by adapting the dynamic range of the feature-contrast response function to the stimulus contrast regime. However, this effect might not

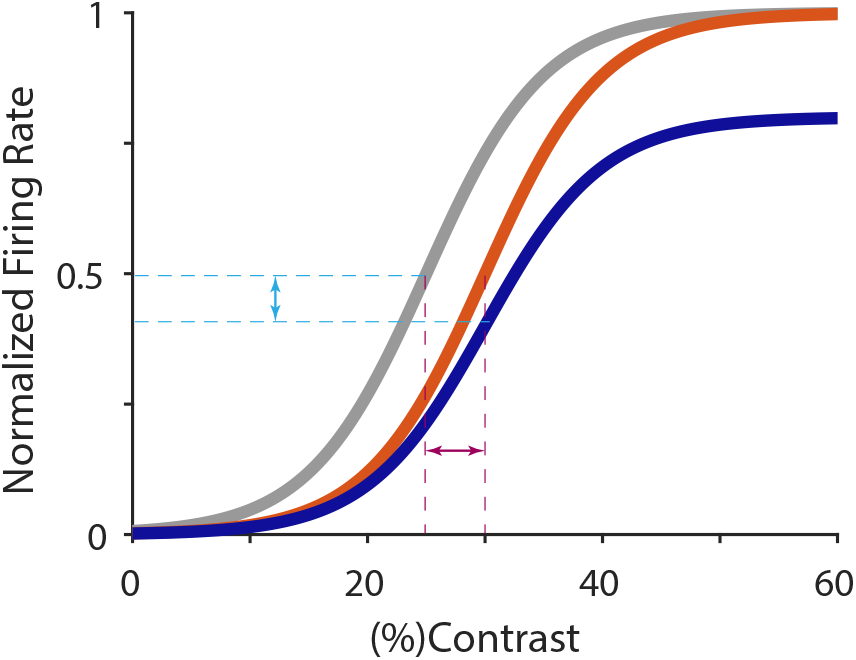
Contrast gain control and response gain control adaptation phenomena, which are often reported for visual neurons. The gray curve illustrates contrast response function that is typical of a V1 cell. In the event of contrast gain control effect, the contrast response function shifts rightward (orange curve), resulting in an increase in the contrast gain (red arrow). In the event of response gain control effect, the contrast response function depresses downward (blue curve), resulting in a decrease in the response gain (cyan arrow).

be significant in some RGCs and, as a result the change in the feature-contrast threshold may not be adequate to improve the sensitivity of these cells to the low-feature contrast of WGN. Hence, some RGCs (or possibly the subsequent cells in the lateral geniculate nucleus) might not be activated during WGN stimulation. These cells would, accordingly, drop out from the feedforward visual stream. The aggregate effect on visual cortical cells could be a reduction in the number of RF filters, as they indicate the effective dimensionality of the cell’s feature space.

### 3.3 Closely related studies

Felsen et al. (2005) measured the sensitivity of cat V1 cells to features in natural and random noise stimuli. While there are similarities between their study and ours, there are also significant differences. (1) They uncovered RFs in cat V1 using NS and spike-triggered average (STA) and spike-triggered covariance (STC) analysis, whereas we used a far more general RF characterization technique (i.e., the NIM), which uses maximum likelihood estimation, thereby, reducing the risk of any bias or artifact in the RF filters. (2) They only used NS stimulation to uncover RF filters, which were then presumed to stay unchanged during any other arbitrary visual stimulation. (3) We used a large stimulus that covered a wide visual field of V1, whereas they used localized stimuli presented only to the RF of each neuron. This means that the neuron’s surrounding cortical areas are most likely in a state that is not very far from the spontaneous state, hence, the effect of lateral connections might not be much affected by the change in the contrast of the stimulus. (4) They compared contrast response functions of V1 cells using NS and random noise images that had equal feature-contrasts. This was done for each cell by creating a random noise sequence that was matched in feature-contrast to the preferred features of the NS for each image frame. In other words, Felsen et al. (2005) studied and compared V1 feature-contrast response functions within the domain of the NS feature-contrast range, whereas we compared the response functions within the space of the WGN feature-contrasts. We did this because we directly stimulated the cells with both WGN and NS. Our findings suggest the emergence of interesting adaptation phenomena, which mean that the RFs behave differently in different feature-contrast regimes. This contradicts the assumption of a fixed set of filters, as used in Felsen et al. (2005) analysis.

Strictly for simple cells in V1, T. O. Sharpee et al. (2006) reported an amplification in the higher spatial frequency (SF) components of the uncovered RF filters when exposed to a change in the statistics of the stimulus from WGN to NS. They interpreted their results to mean that the visual system was compensating for the under-represented higher SF components in NS,

to improve the efficiency of neuronal information transmission. However, the mere adjustment of amplitude spectra of RF filters in the visual system cannot fully account for all the statistical changes in the visual input to the system, because only second-order statistics of the stimulus ensemble are reflected by the amplitude spectrum (see also Felsen et al. (2005)). Rather, the spatial (Fourier) phase spectrum of image ensembles mainly captures information corresponding to higher-order statistics in the image that describe features such as edges, contours, curves etc. Our results corroborate T. O. Sharpee et al. ‘s findings; that is with NS stimulation, V1 cells tend to have RF filters with higher SF preferences and narrower SF bandwidths (Fig. 9). However, our findings extend beyond simple cells as we recorded from cells with multiple filters. Our data unveil increased numbers of filters and changed feature-contrast response functions as well as increased SF tuning. These modifications optimize the dynamic ranges of most recorded V1 cells for the given image statistics. The more narrowly SF tuned RF filters in response to NS stimuli imply a redundancy reduction in the input representation using less-active units, consistent with the efficient coding hypothesis.

The bias towards RF filters with lower SF preferences and broader SF bandwidths identified using WGN might be explained also by the effects that contrast adaptation imposes on the visual system. The effects of contrast gain control for ON cells are more pronounced than the effects for OFF cells in the early visual pathway (Chander & Chichilnisky, 2001; Felsen et al., 2005; Ratliff, Borghuis, Kao, Sterling, & Balasubramanian, 2010; Zaghloul et al., 2005). When switching from high to low contrast regimes, the contrast response functions of OFF cells show little adaptation to changes in contrast, thereby the reduction in their contrast gain function (i.e., the shift to low contrast) is less pronounced compared to ON cells (Bonin et al., 2006). Furthermore, OFF cells are more selective to high SF features (perhaps due to their small dendritic fields) in the visual scene than ON cells (Ratliff et al., 2010; also see Chichilnisky & Kalmar, 2002). The less pronounced contrast adaptation effect in OFF cells indicates that, when switching from high to low contrast stimulation (e.g., from NS to WGN), some OFF cells likely exhibit little contrast gain control. Therefore, the reduction in their contrast gain is not sufficient to improve their sensitivity to the low contrast stimulus. As these cells all feed into visual cortical cells, the reduction in the activity of such OFF cells would propagate in the feedforward stream of the visual system. Therefore, the reduced OFF-cell input might explain the significant reduction in the SF preferences and broadening of SF bandwidths of cortical RF filters during WGN stimulation, because OFF cells tend to be selective for high SFs.

### 3.4 Function

In most V1 cells in our study, the changes in the RFs between the WGN and NS regimes can be interpreted as a mechanism to increase the amount of information encoded by the cell. Cells adapt their feature-contrast response functions to the stimulus’ dynamic range, and this change seems to be related to the range of feature-contrasts in the stimulus. There are changes that appear in the shape of the response functions. Higher response gains are allocated to the stimuli that are rare in the image, which belong to both the positive and negative tails of the feature-contrast distributions. Such an encoding mechanism carries more information about the stimulus (Felsen et al., 2005).

The changes in feature-contrast response functions observed in our V1 data are consistent with the contrast adaptation phenomena reported in the primary visual cortex, described by contrast gain control or response gain control effects (Albrecht et al., 1984; Ohzawa et al., 1985). When the stimulus contrast changes from low to high, in the event of a contrast gain control effect, the response function to high contrast stimuli is shifted rightward to higher contrasts (Fig. 10; gray vs. orange curves), while with response gain control effect the response function is shifted downward to lower spike rates (Fig. 10; gray vs. blue curves). Both effects result in a decrease in the response function to higher contrast stimuli compared to the low contrast response function.

The findings in the present study can have implications for the mechanisms employed by the visual system to deal with the uncertainty of visual information. In retina, OFF cells have smaller dendritic field sizes than ON cells, but they outnumber the ON cells two-fold. In the low feature-contrast regime (e.g., WGN), the uncertainty in the visual stimulus is higher than in the high feature-contrast regime, due to the signal being inherently noisy. In this case, the visual system can improve its performance by increasing its sensitivity to coarser features in the visual input. This can be done by shutting down some of the high spatial frequency filtering channels in the early visual processing stages. The visual system could implement this via adaptive contrast gain control mechanisms in OFF cells in early processing stages, or through cortical processing by a change in the balance between feedforward and lateral connections due to the change in the stimulus contrast as conjectured by (Nauhaus, Busse, Carandini, & Ringach, 2009).

Overall, our findings conclude that the primary visual cortex extracts image features from the input in a plastic way, which is formed based on the nature of the stimulus.

## 4 Methods

### 4.1 Preparation and surgery

Extracellular recordings were made primarily from Area 17 in cat cortex, but some recording locations were on the border with Area 18, making unequivocal identification difficult. As both areas are retinorecipient, we refer to our recordings as being from primary visual cortex (V1). Recordings were made from V1 in six anesthetized cats using methods described previously (Meffin et al. 2015; Almasi et al. 2020; Sun et al. 2021). Experiments were conducted according to the National Health and Medical Research Council’s Australian Code of Practice for the Care and Use of Animals for Scientific Purposes. All experimental procedures were approved by the Animal Care Ethics Committee at the University of Melbourne (ethics ID 1413312).

Anesthesia was induced in adult cats (2-6 kg) with an intramuscular injection of ketamine hydrochloride (20 mg/kg i.m.) and xylazine (1 mg/kg). The cats were intubated, cannulated, and placed in a stereotaxic frame. Once intubated, oxygen and isoflurane (1-2%) were used to maintain deep anesthesia during all surgical procedures. A craniotomy was performed to expose cortical areas 17 and 18. Isoflurane was used during the surgery because it is safe for humans. Anesthesia was switched to gaseous halothane in a fully closed system during data recording (0.5-0.7%) and the depth of anesthesia was determined by monitoring a variety of standard indicators (see Sun et al. 2021). Halothane was used during recordings because it has been shown to maintain anaesthesia but have a less suppressive impact on cortical responses (Villeneuve and Casanova 2003). To avoid eye movements during recordings, muscular blockade was induced and maintained with an intravenous infusion of vecuronium bromide at a rate of 0.1 mg·kg^−1^·h^−1^.

Mechanical ventilation was utilized to maintain end-tidal CO_2_ between 3.5% and 4.5%. After an experiment the animal was humanely killed without regaining consciousness with an intravenous injection of an overdose of barbiturate (pentobarbital sodium, 150 mg/kg). Animals were then perfused immediately through the left ventricle of the heart with 0.9% saline followed by 10% formol saline, and the brain extracted.

### 4.2 Visual stimuli and data recording

Visual stimuli were generated using a ViSaGe visual stimulus generator (Cambridge Research System, Cambridge, UK) on a calibrated, Gamma corrected LCD monitor (ASUS VG248QE, 1920×1080 pixels, refresh rate 60 Hz, 1 ms response time) at a viewing distance of 57 cm. White Gaussian noise (WGN) and Natural Scene (NS) stimuli comprising 90×90 pixels over 30*°* of the visual field were employed to estimate the neuronal receptive fields of V1 cells using the NIM framework (see Methods below). The WGN and NS images used in the stimuli had a mean value equal to the mid-luminance of the display monitor. The WGN images had a standard deviation chosen to result in a 10% saturation rate for individual pixels, i.e. the mean had a normalized intensity of 0.5, and 10% of pixels had a value of either 0 or 1 corresponding to the lowest and highest luminance of the monitor. The NS stimuli comprised 90×90 pixels and were randomly extracted image patches from a database of natural images (Van Hateren and Van der Schaaf 1998). Each NS stimulus block contained patches that were drawn from 100 randomly chosen images in the database, with each image 1536×1024 pixels. Both WGN and NS stimuli had their global RMS contrast matched, which was set to ∼0.3. WGN and NS stimuli were presented in separate blocks of 12,000 images and with blocks interleaved to ensure the physiological comparability of the recordings. Each image frame was presented for 1/30 s, followed by a blank screen of the mean luminance (intensity = 0.5), displayed for the same duration in blocks of 12,000. The blank period aimed to increase the overall response of the cell to the stimuli by increasing the temporal contrast. The total duration of a block was ∼14 min.

Extracellular recordings were made with single shank probes with Iridium electrodes (linear 32-electrode arrays, 6 mm length, 100 *µ*m electrode site spacing; NeuroNexus), which were inserted vertically using a piezoelectric drive (Burleigh inchworm and 6000 controller, Burleigh instruments, Rochester, NY). Extracellular signals were acquired from 32 channels simultaneously using a CerePlex acquisition system and Central software (Blackrock Microsystems, Salt Lake City, Utah) sampled at 30 kHz and 16-Bit resolution on each channel. Filtering was performed by post-processing.

### 4.3 Post-processing and spike sorting

Spike sorting of recordings was performed using KiloSort (Pachitariu, Steinmetz, Kadir, Carandini, & Harris, 2016) and the graphical user interface *phy* (Rossant et al., 2016). Single units were identified as previously described in Almasi et al. (2020).

### 4.4 Model estimation

#### 4.4.1 Model definition and parameters estimation

We have employed an adapted version (Almasi et al., 2020) of the nonlinear input model (NIM), originally introduced by McFarland et al. (2013). The model is depicted in Figure 1a and describes

the firing rate of the cell as a nonlinear function of the input stimulus,

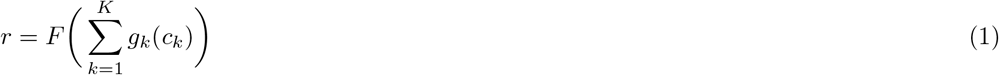

where *c*_*k*_ = **h**_*k*_ · **s** is termed the feature-contrast of the stimulus **s** with respect to the spatial filter **h**_*k*_, which is defined as their inner product. The model cell conceptually sums inputs from *K* parallel synaptic input streams, which are determined as a hyperparameter of the model, to give a generator potential 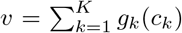. Each input is determined by an arbitrary function *g*_*k*_(·) of the feature-contrast *c*_*k*_ of filter **h**_*k*_, which is called the input function and captures the processing performed by one or more presynaptic neurons. The number of input streams *K* (i.e., RF filters) for each cell is determined using a statistical significance test described below. The function *F* (·) gives the overall spiking nonlinearity of the cell that converts the generator potential into firing rates and is described using a parametric representation as

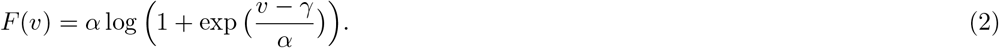

The model assumes that the responses *R*_*obs*_ = {*R*^(1)^, …, *R*^(*T*)^} (integer spike counts) to the presented set of mutually independent stimuli *S* = {**s**^(1)^, …, **s**^(*T*)^} follow a homogenous Poisson distribution function

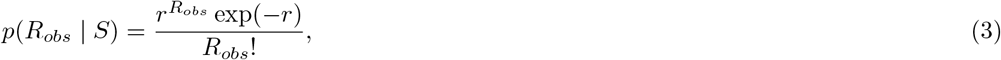

where *r* is the firing rate function described in Eq. 1. The spike counts for each stimulus were obtained by pooling all the spikes that occurred after the stimulus presentation within a window that had a duration equal to the presentation period (1/30 s) shifted by a certain latency. The latency was estimated for each cell based on post-stimulus time histograms (PSTH), which clearly revealed the time of response onset, to maximize the mean response to the NS and WGN. The estimation of the response latency was achieved by fitting a von Mises function to the PSTH of the cell’s response to each stimulus block, and later by averaging across different recordings to give the estimate. The estimation of the model parameters (spatial filters, input functions and spiking nonlinearity) was obtained simultaneously by maximizing the (log)-likelihood of the model given the stimuli and their evoked responses. The reader is advised to refer to Almasi et al. (2020) for comprehensive details about the model and the simultaneous estimation procedure.

#### 4.4.2 Significance test to determine the number of RF filters

We determined the number of spatial filters within the receptive field of each cell using cross-validation. In doing so, the number of filters for each cell was systematically varied while the statistical significance of each filter was evaluated by bootstrapping. For this, we divided the data into a training set, which comprised four-fifths of the data, and a test set, which comprised the other one-fifth of the data. For each specified number of filters, we used the training set to estimate the filters, and then assessed the performance of the model by computing its log-likelihood using resampling from the test set (this was repeated 500 times). Thus, for each number of filters, we found a distribution for the log-likelihood computed on the test set. The inclusion of a new filter was counted as significant if it significantly improved the log-likelihood of the model on the test set (Z-score *>* 2). Technical details of implementing this test is given in Almasi et al. (2020).

### 4.5 Estimation of response latency and strength

The latency in the response to the stimulus was obtained by fitting a *von Mises* distribution defined as

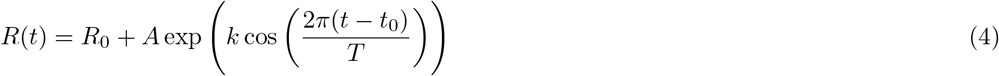

to the response PSTH, for each stimulus recording. In above, *T* = 67 ms and *R*(*t*) represents the response PSTH, *t* ∈ [0, *T*] which indicates the presentation time of a stimulus image followed by a blank image each for 1/30 s. A least-square-error fit of the above curve to the response PSTH was found to identify the parameters *R*_0_, *A, k* and *t*_0_. The curve fitting was performed by the *lsqnonlin* function in Matlab©. The latency, defined as the onset in the response, is calculated to be *t*_0_ − *T/*4. The parameters of the fitted curve are obtained by minimizing the mean squared error between the fitted and actual response PSTH of each stimulus recording. The quality of the fitted curves was assessed using a *r*^2^ goodness-of-fit measure. Fits with *r*^2^ *<* 0.8 were excluded from the latency and spike rate analysis. Overall, the *r*^2^ of the fits formed a distribution with a mean of 0.96 and a median of 0.97. The response latency was averaged across different stimulus presentations to account for change in the response latency due to any change in the neuronal physiological states. Figure A.1 shows the (normalized) fitted PSTH response curves for NS and WGN stimuli, for which the latency in the response was accounted. In this case, the onset of the response is aligned with the stimulus onset. The response strength is defined as the maximum firing rate as impulse per second (ips) in the response PSTH to each stimulus, averaged across different stimulus blocks.

### 4.6 Feature-contrast

The output of the linear filtering stage in the model indicates the similarity between the visual input and the spatial structure of the filter. The authors have previously demonstrated that the filter’s output can be described in terms of RMS or Michelson contrast of the spatial structure (the feature) of that filter embedded in the visual stimulus, hence defined as the feature-contrast (Almasi et al., 2020). Furthermore, the feature-contrast corresponding to a cell’s RF filter can be interpreted as the local contrast of the stimulus when projected onto that filter.

### 4.7 Feature-contrast range

Feature contrast is defined as the output of the spatial RF filters. If the cell has only one RF filter uncovered by WGN, the RF filters’ feature-contrast on WGN follows a univariate Normal distribution with standard deviation of *σ*^*wn*^. In this case, the range of feature-contrast on WGN stimuli is measured as Σ_*wgn*_ = [−3*σ*^*wn*^, 3*σ*^*wn*^]. If the cell has two RF filters uncovered by WGN, their feature-contrast on WGN follows a bivariate Normal distribution. Since there is no orthogonality constraint on RF filters in the NIM, this distribution can in general have an elliptical (correlational) form. We decorrelated this distribution by performing singular value decomposition analysis and obtained a canonical distribution. We then found standard deviations of this distribution along its two canonical axes 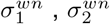. These two values are not necessarily the same, but in our experience, they were found to be very close. We define the range of feature-contrast on WGN as Σ_*wgn*_ = [−3*σ*^*wn*^, 3*σ*^*wn*^] where *Σ*^*wn*^ is 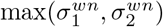. This measure can be easily generalized to WGN RF dimensions greater than two, wherein the WGN feature-contrast follows a multivariate Normal distribution.

To find the distribution of the feature-contrast for NS RF filters when using NS stimuli, we performed the same procedure explained above. After obtaining the canonical (decorrelated) distribution of NS feature-contrast, we found NS stimuli that lie inside a ball (or hyper-ball) with a radius equal to Σ_*wgn*_. This subset of NS stimuli is referred to as matched-feature-contrast-NS stimuli, and results in a distribution of feature-contrast for NS RF filters that are within the range of feature-contrasts of the WGN RF filters on WGN stimuli.

### 4.8 Neuron’s feature space

The RF filters of a neuron identified using the NIM framework will span a space that is termed the neuron’s feature space. Mathematically, this feature space is equal to the column space (ℍ) of the matrix collecting all the RF filters as its columns *H* = [**h**_1_ … **h**_*m*_], where *m* denotes the number of RF filters. The feature space by definition contains all possible linear combinations of the neuron’s RF filters:

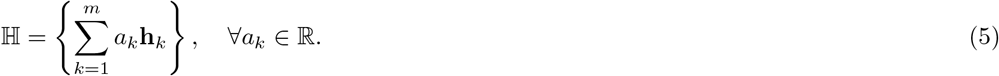

### 4.9 Control analysis to match spike count

This analysis was performed for the cells that met three criteria: (1) they responded with a higher spike-rate to NS than to WGN stimuli, (2) they had their RF uncovered using both WGN and NS, and (3) they had more RF filters uncovered using NS than using WGN stimuli. This control analysis corrects for the higher spike-rates on NS, by matching the NS spike rate to the spike-rate recorded on WGN stimuli. This was done by randomly sampling spikes from the cell’s spike train in response to NS stimuli until the specified spike count for WGN was reached.

## A Comparison of predictive ability of models with WGN and NS feature spaces on WGN data

We asked whether the greater number of RF filters uncovered by NS for most cells are the fundamental set of features for cells in order to operate well under any stimulation regime. To evaluate the above hypothesis, we first formed and trained the following models on a WGN training set:

1. The model whose RF filters are held fixed to the WGN RF filters projected onto the NS feature space, denoted by 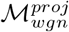.
2. The model whose RF filters are held fixed to the NS RF filters. We refer to this model as the feature-extended-WGN-NIM, denoted by 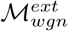.

In above, the latter model 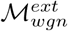 incorporates all the feature dimensions that amount to ℍ_*ns*_, hypothesized necessary for the model to perform well under any stimulus regime. We fitted both models using a WGN training set, by estimating the RF nonlinearities, while the RF filters were held fixed. If, as hypothesized, ℍ_*ns*_ is the true feature space of the cell, then 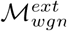 must outperform 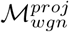 when they are compared against a held-out WGN test set (note that 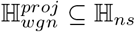).

Our analysis showed that for a majority (80%) of cells in our population of V1 data, 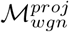 significantly outperformed 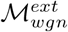 (unpaired t-test; *p <* 0.001). For a few minority (20%) of cells, we found no significant difference between the performance of the two models, namely both performed equally on the hold-out WGN test data. It may seem odd that 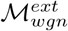 can actually perform worse than 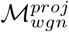 that has less features. Given that 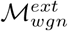 contains additional features with meaningful spatial structures, it may seem logical that it has to perform at least as good as a model with a subset of its RF filters. However, it should be noted that due to the greater number of filters in the model, it is possible that the model overfits to the training data, and when assessed against the hold-out test data, it does not generalize to the test set as good as a simpler model with less RF filters like 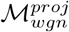.

**Figure A.1.**
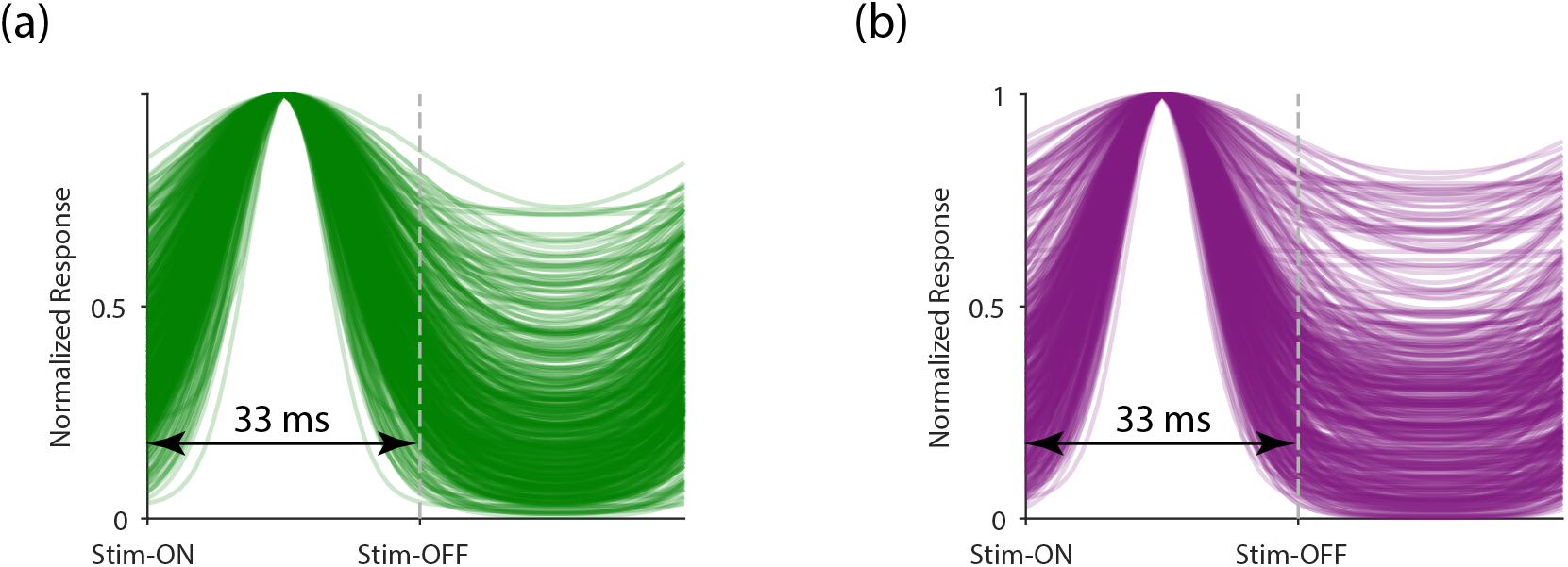
Post-Stimulus Time Histogram (PSTH) of the responses of all the units analyzed in this study, overlaid in a single plot for (**a**) NS and (**b**) WGN stimulation. Each graph is obtained by correcting for the latency in the response. Graphs are aligned with the time onset of the stimulus (Stim-ON). These plots confirm that the peak responses occurred within the 33ms window, which was used to count spikes for RF estimation using the NIM.

**Figure A.2.**
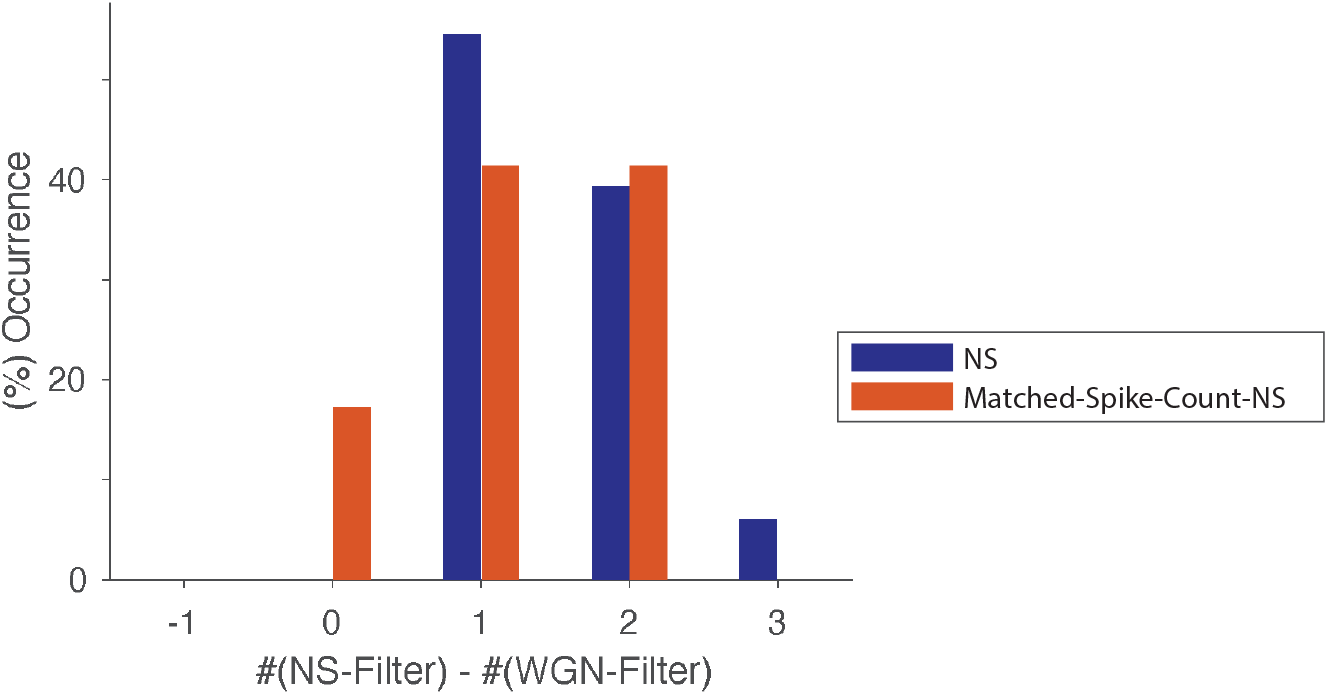
Bar graphs show the difference between the number of RF filters uncovered using NS and WGN for the cells that had more filters using NS and higher spike counts to NS (dark-blue), and for the same cells when their spike count to NS was matched to that to WGN stimuli (orange).

## Notes

### Competing Interest Statement

The authors have declared no competing interest.

